# CarrierSeq: a sequence analysis workflow for low-input nanopore sequencing

**DOI:** 10.1101/175281

**Authors:** Angel Mojarro, Julie Hachey, Gary Ruvkun, Maria T. Zuber, Christopher E. Carr

## Abstract

**Motivation:** Long-read nanopore sequencing technology is of particular significance for taxonomic identification at or below the species level. For many environmental samples, the total extractable DNA is far below the current input requirements of nanopore sequencing, preventing “sample to sequence” metagenomics from low-biomass or recalcitrant samples.

**Results:** Here we address this problem by employing carrier sequencing, a method to sequence low-input DNA by preparing the target DNA with a genomic carrier to achieve ideal library preparation and sequencing stoichiometry without amplification. We then use CarrierSeq, a sequence analysis workflow to identify the low-input target reads from the genomic carrier. We tested CarrierSeq experimentally by sequencing from a combination of 0.2 ng *Bacillus subtilis* ATCC 6633 DNA in a background of 1 μg *Enterobacteria phage λ* DNA. After filtering of carrier, low quality, and low complexity reads, we detected target reads (*B. subtilis*), contamination reads, and “high quality noise reads” (HQNRs) not mapping to the carrier, target or known lab contaminants. These reads appear to be artifacts of the nanopore sequencing process as they are associated with specific channels (pores). By treating reads as a Poisson arrival process, we implement a statistical test to reject data from channels dominated by HQNRs while retaining target reads.

**Availability:** CarrierSeq is an open-source bash script with supporting python scripts which leverage a variety of bioinformatics software packages on macOS and Ubuntu. Supplemental documentation is available from Github - https://github.com/amojarro/carrierseq. In addition, we have compiled all required dependencies in a Docker image available from - https://hub.docker.com/r/mojarro/carrierseq.

## 1 Introduction

Environmental metagenomic sequencing poses a number of challenges. First, complex soil matrices and tough-to-lyse organisms can frustrate the extraction of deoxyribonucleic acid (DNA) and ribonucleic acid (RNA) (Lever et al., 2015). Second, low-biomass samples require further extraction and concentration steps which increase the likelihood of contamination (Barton et al., 2006). Third, whole genome amplification may bias population results (Sabina and Leamon, 2015) while targeted amplification (e.g., 16S rRNA amplicon) may decrease taxonomic resolution (Poretsky et al., 2014). To address these challenges, we have developed extraction protocols compatible with low-biomass recalcitrant samples and difficult to lyse organisms (Mojarro, Ruvkun, et al., 2017). These protocols, developed using tough-to-lyse spores of Bacillus subtilis, allow us to achieve at least 5% extraction yield from a 50 mg sample containing 2 x 105 cells/g of soil without centrifugation (Carr et al., 2017). Furthermore, in order to avoid possible amplification biases and additional points of contamination, we have experimented with utilizing a genomic carrier (*Enterobacteria phage λ*) to shuttle low-input amounts of target DNA (*B. subtilis*) through library preparation and sequencing with ideal stoichiometry (Mojarro, Hachey, et al., 2017). This approach has allowed us to detect down to 0.2 ng of *B. subtilis* DNA prepared with 1 μg of Lambda DNA using the Oxford Nanopore Technologies (ONT) MinION sequencer (supplementary data, https://www.ncbi.nlm.nih.gov/bioproject/398368). Here we present CarrierSeq, a sequence analysis workflow developed to identify target reads from a low-input sequencing run employing a genomic carrier.

## 2 Methods

CarrierSeq implements bwa-mem (Li, 2013) to first map all reads to the genomic carrier then extracts unmapped reads by using samtools (Li et al., 2009) and seqtk (Li, 2012). Thereafter, the user can define a quality score threshold and CarrierSeq proceeds to discard low-complexity reads (Morgulis et al., 2006) with fqtrim (Pertea, 2015). This set of unmapped and filtered reads are labeled “reads of interest” (ROI) and should theoretically comprise target reads and likely contamination. However, ROIs also include “high-quality noise reads” (HQNRs), defined as reads that satisfy quality score and complexity filters yet do not match to any database and disproportionately originate from specific channels. By treating reads as a Poisson arrival process, CarrierSeq models the expected ROIs channel distribution and rejects data from channels exceeding a reads/channels threshold (x_crit_) (Figure 1).

**Fig. 1.**
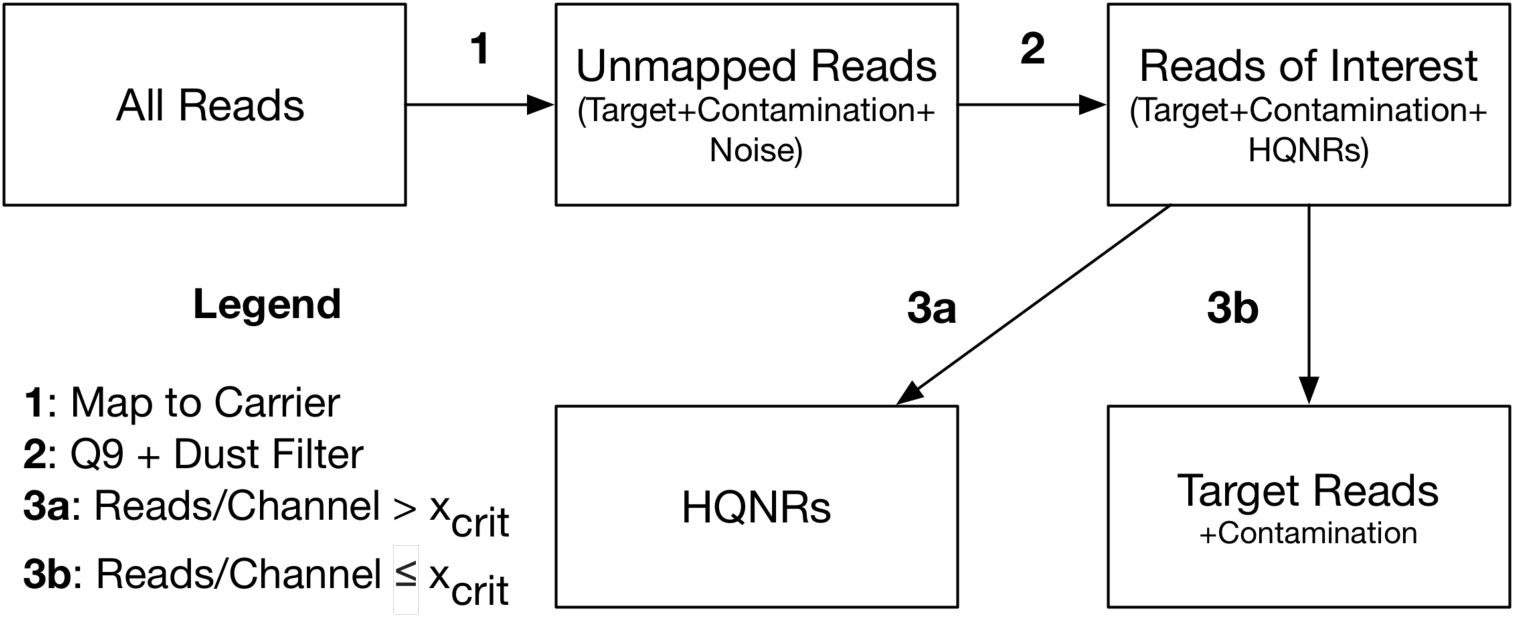
CarrierSeq workflow. Starting from all reads, CarrierSeq identifies unmapped reads then applies a quality score and complexity filter to discard low-quality reads. Afterwards, CarrierSeq applies a Poisson distribution test to sort *likely* high-quality noise reads (HQNRs) from target reads.

### 2.1 Quality Score Filter

The default per-read quality score threshold (Q9) was determined through receiver operating characteristic curve (ROC) analysis (Fawcett, 2006) of carrier sequencing runs of *B. subtilis* and Lambda DNA (Figure 2). This threshold is best suited for Lambda carriers that are 99% library by mass and essentially function as a pseudo “lambda burn-in” experiment (Nanoporetech.com, 2017). Therefore, the user is encouraged to define their own threshold based on their libraries’ quality control metrics (e.g., carrier to target ratio, quality distribution, sequencing accuracy achieved, and basecaller confidence).

**Fig. 2.**
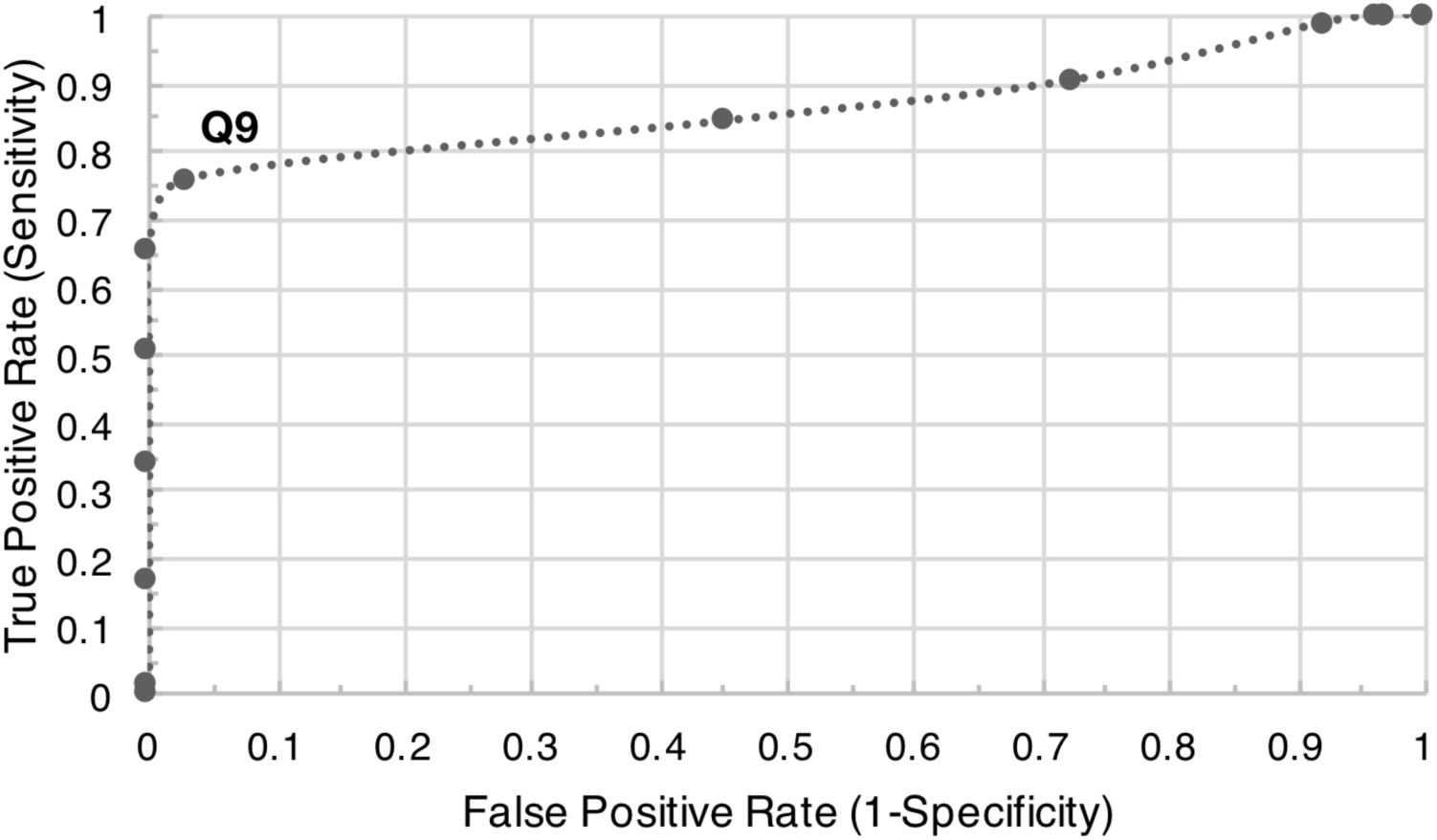
Receiver operating characteristic curve. Q9 provides a good threshold which discards the majority of low-quality and noise reads (0.76 True Positive Rate and 0.03 False Positive Rate) for carrier runs that are 99% Lambda DNA by mass. A perfect quality score threshold would plot in the top left of the ROC curve.

### 2.2 Poisson Distribution Sorting

Assuming that sequencing is a stochastic process, CarrierSeq is able to identify channels producing spurious reads by calculating the expected Poisson distribution of reads/channel. Given total ROIs and number of active sequencing channels, CarrierSeq will determine the arrival rate (λ = reads of interest/active channels). CarrierSeq then calculates an x_crit_ threshold (x_crit_ = poisson.ppf (1 – p-value), λ)) and sorts ROIs into target reads (reads/channel ≤ x_crit_) or HQNRs (reads/channel > x_crit_) (supplementary data).

### 2.3 Implementation

Reads to be analyzed must be compiled into a single fastq file and the carrier reference genome must be in fasta format. Run CarrierSeq with:

`./carrierseq.sh –i <input.fastq> –r <reference.fasta> –o <output_directory>`

## 3 Results & Discussion

From experimenting with low-input carrier sequencing and CarrierSeq we observed that the abundance of HQNRs may vary per run, perhaps due to sub-optimal library preparation, delays in initializing sequencing, or other sequencing conditions. In addition, target DNA purity and lysis carryover (e.g., proteins) may conceivably contribute to HQNR abundance. Possibly due to pore blockages from unknown macromolecules that result in erroneous reads. While the cause or significance of HQNRs have yet to be determined, future work will focus on developing a method to identify HQNRs on a per-read basis. In contrast, the current approach discards entire HQNR-associated channels at the risk of discarding target reads. Moreover, some reads in non-HQNR-associated channels may also be artifacts. The ability to identify HQNRs on a per-read basis is especially important for metagenomic studies of novel microbial communities where HQNRs may complicate the identification of an unknown organism, or in a life detection application (Carr et al., 2017) where artefactual reads not mapping to known life could represent a false-positive.

## 4 Summary

CarrierSeq was developed to analyze low-input carrier sequencing data and identify target reads. We have since deployed CarrierSeq to test the limits of detection of ONT’s MinION sequencer from 0.2 ng down to 2 pg of low-input carrier sequencing. CarrierSeq may be a particularly valuable tool for in-situ metagenomic studies where limited sample availability (e.g., low biomass environmental samples) and laboratory resources (i.e., field deployments) may benefit from sequencing with a genomic carrier.

## Acknowledgements

The authors would like to thank Michael Micorescu at Oxford Nanopore Technologies for providing and granting us permission to utilize his fastq quality filter script.

